# Two decades of suspect evidence for adaptive DNA-sequence evolution - Failure in consistent detection of positive selection

**DOI:** 10.1101/417717

**Authors:** Ziwen He, Qipian Chen, Hao Yang, Qingjian Chen, Suhua Shi, Chung-I Wu

## Abstract

A recent study suggests that the evidence of adaptive DNA sequence evolution accumulated in the last 20 years may be suspect^1^. The suspicion thus calls for a re-examination of the reported evidence. The two main lines of evidence are from the McDonald-Kreitman (MK) test, which compares divergence and polymorphism data, and the PAML test, which analyzes multi-species divergence data. Here, we apply these two tests concurrently on the genomic data of *Drosophila* and *Arabidopsis*. To our surprise, the >100 genes identified by the two tests do not overlap beyond random expectations. The results could mean i) high false positives by either test or ii) high false-negatives by both tests due to low powers. To rule out the latter, we merge every 20 - 30 genes into a “supergene”. At the supergene level, the power of detection is high, with 8% - 56% yielding adaptive signals. Nevertheless, the calls still do not overlap. Since it is unlikely that one test is largely correct and the other is mostly wrong (see Discussion), the total evidence of adaptive DNA sequence evolution should be deemed unreliable. As suggested by Chen *et al.*^1^, the reported evidence for positive selection may in fact be signals of fluctuating negative selection, which are handled differently by the two tests. Possible paths forward on this central evolutionary issue are discussed.

## Introduction

The inferences of adaptive evolution in DNA sequences permit the assessment of the biological significance of genes of interest. Such inferences may then guide the planning of functional validation. Extensive reports of adaptively evolving genes can be found in almost all taxa^1–4^ as well as all types of cancers^5,6^. Indeed, the large-scale genomic data amassed in the last two decades have led to the acceptance of pervasive adaptive evolution over the neutral theory of molecular evolution^4,7–10^.

In a companion study, we question this acceptance because positive selection often yields the same signals as reduced negative selection, for example, due to decreased population sizes. Nevertheless, plausible skepticisms are not grounds for rejection. It is necessary to re-examine the analyses that reported adaptive signals in DNA sequences. The detection of positive selection largely falls into two broad classes^11–15^. One class attempts to detect positive selection that operates within populations^11,14,16^. The other focuses on positive selection that operates in the longer term, i.e., the divergence between species^17–19^. Methods of either class may use data of both polymorphism and divergence^13,17,20^. Positive selection signals could be abundant between species but undetectable within populations, or vice versa. It is hence possible to reject the neutral theory in part (either within or between species) or in whole.

In this study, we focus on the between-species tests. In such tests, one compares the number of non-synonymous changes per non-synonymous site (Ka or dN) vs. the per-site synonymous changes (Ks or dS)^12,18,21^. The Ka/Ks (or dN/dS) ratio would deviate from 1 if nonsynonymous changes are under stronger selection than synonymous substitutions. In the absence of selection, R = Ka/Ks ~ 1, which is the hallmark of neutral evolution^22,23^. In among-species comparisons, genome-wide R ranges mainly between 0.05 and 0.25^23,24^, thus indicating the prevalence of negative selection. When R > 1, positive selection is evident. However, R > 1 is too stringent a criterion as it requires positive selection to overwhelm negative selection. Indeed, few genes in any genome comparison have R significantly greater than 1^15,25^.

The two commonly used methods that relax the requirement for R > 1 over the entire gene are the MK (McDonald-Kreitman)^13,17^ and the PAML (Phylogenetic analysis by maximum likelihood)^26,27^ tests. Each test relies on a different set of assumptions that cannot be easily verified. For that reason, it remains a serious concern that either or even both tests may yield a large fraction of false positives. If the detected adaptive signals are true, the results from the two tests are expected to show substantial overlap (see Discussion for details).

All tests need to heed the caveat of low power in detecting selection due to the small number of changes in each gene. If both tests have low powers (and hence high false negatives), the overlap would be low, even if both tests have correctly identified adaptive genes with few false positives. For that reason, pooling sites would be necessary. After all, even single genes represent pools of sites of various adaptive values and previous studies have often combined sites from the whole genome to raise the statistical power^3,4,20^. In this study, we would raise the power of detecting selection by pooling genes in different combinations.

## Theoretical background

While Ka and Ks are the cornerstones for detecting natural selection, they can only inform about either positive or negative selection, but not both. This is because Ka/Ks, when averaged over all sites, is the joint outcome of the two opposing forces, as outline below using the basic population genetic theory:

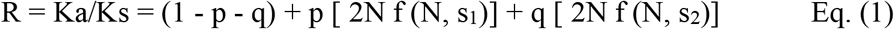

where p and q are the proportion of advantageous and deleterious mutations, respectively^22,28,29^; f (N, s) = (1 − e^−s^)/ (1 − e^−2Ns^) is the fixation probability of a mutation with a selective coefficient s that can be > 0 (denoted by s_1_) or < 0 (s_2_) and half the value in heterozygotes^29,30^ (See Supplementary Information for details). N is the effective population size. For example, if Ka/Ks = 0.2, then the null hypothesis is the neutrality with q = 0.8, p = 0 and f (N, s_2_) = 0 (i.e., no fixation of deleterious mutations). The alternative hypothesis would be adaptive evolution with p > 0 and f (N, s_1_) > 0 (fixation of advantageous mutations).

To tease apart positive and negative selection, one often uses DNA sequences from several closely-related species, some of which should have polymorphism data. For this study, the data are from species in the *Drosophila* and *Arabidopsis* clade, respectively (Fig. 1). The hypothesis testing for positive selection by the MK test is done on a particular phylogenetic lineage, marked by red line in Fig. 1. The Ka and Ks values in the red line lineage are contrasted with the corresponding polymorphisms (Pa and Ps) in the blue triangle. The value of Pa and Ps denotes, respectively, the level of nonsynonymous and synonymous polymorphism (per site) within a species. The rationale of the MK test is that p ~ 0 in the polymorphism data thanks to the rapidity with which advantageous mutations are fixed. Thus, Eq. (1) becomes

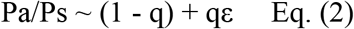

where ε represents the amount of deleterious polymorphism and should be a very small number. In short, the MK test estimates q from Eq. (2) and then extracts p from Eq. (1).

**Figure 1.**
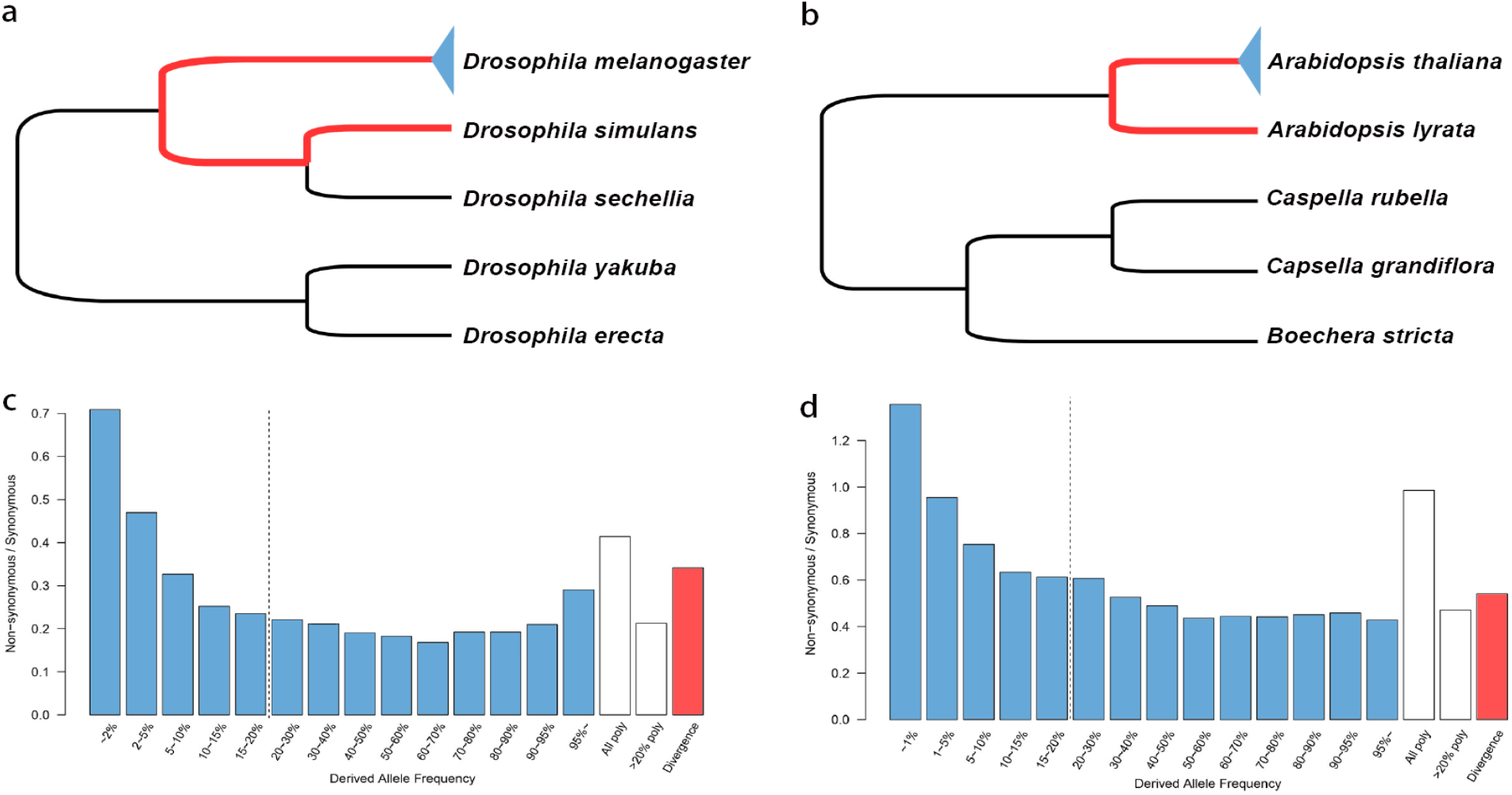
Between-species divergence and within-species polymorphism for detecting positive selection. **a** and **b**, Phylogeny of *Drosophila* and *Arabidopsis* species. Both the MK and PAML tests are forced to detect positive selection along the branches marked by red. The MK test uses polymorphisms (indicated by the blue triangle) for reference. The reference for PAML is described in Methods. **c** and **d**, The A/S ratio as a function of the mutant frequency in *D. melanogaster* and *A. thaliana*, where A is non-synonymous, and S is synonymous polymorphism. The dashed line, separating low- and high-frequency bins, is placed where the A/S ratio reaches a steady level. Open bars on the right are, respectively, A/S ratios for all bins and for high-frequency bins. The divergence A/S ratio is shown as the red bar.

There are, however, several difficulties in interpreting the MK test results. First, the strength of negative selection is estimated from the recent evolutionary history (the blue triangle in Fig. 1), whereas positive selection is inferred from a different lineage (the red line). As pointed out before, an increase in the effective size of the extant population would lead to the under-estimation of Pa/Ps and, thus, an over-estimation of positive selection^31,32^. Second, the estimation of negative selection is not straightforward. The Pa/Ps ratio would decrease as the variant frequency increases and may increase again when the mutant frequency approaches 1^4,5,31,33^. Both patterns can be seen in *Drosophila* (Fig. 1c). In *Arabidopsis* (Fig. 1d), the pattern is similar at the low frequency end, but not at high frequencies. Given the complex patterns, accurate estimation of negative selection is not straightforward in the MK test^2–4,13,17,31,32,34,35^. Third, the MK test is strictly applicable only to sites that share the same genealogy. In the presence of recombination, in particular when unlinked loci are used, biases could be non-trivial, making corrections necessary^3,4^.

The other widely used approach to estimating adaptive evolution is the PAML method^26,27^. PAML compares the substitution numbers across many lineages to identify positively (or negatively) selected genes on the assumption that unusually high (or low) numbers could be indicative of selection. In particular, the proportion of adaptive sites that have a higher non-synonymous than neutral rate is estimated by PAML. There are three (sub-) models in PAML, each representing a different set of assumptions. The site model identifies sites with an increase or decrease in non-synonymous substitutions in the entire phylogeny^19,26,27^. The branch-site model compares sites of a pre-selected branch (the foreground) to other sites on all branches as well as the same sites on other branches (the background)^9,36,37^. The third sub-model is not considered here.

Despite the very different approaches, the MK and PAML tests can be used to answer the same question – How much adaptive evolution has happened in the chosen genes on a given branch (e.g., the red-line branch of Fig. 1a and 1b)? Because the MK test is about positive selection along the red line, it does not offer any information about selection elsewhere in the phylogeny. Therefore, it is necessary to compare it to each of the two PAML sub-models. If the MK test identifies genes that are generally prone to adaptive evolution, the proper comparison would be the PAML site model. Alternatively, if the adaptation is specific to a specific branch, then the branch-site model would be a more suitable comparison. We will present the site model results in the main text and the branch-site model results in Supplementary Information. The two sets of comparisons lead to the same conclusion although the site model appears to be statistically more robust.

In short, this study follows the well-known properties of MK and PAML, obtained from theories and extensive simulations^17,19,20,32,38^ to their logical conclusions. Since no new theory was developed here, the discordance between the two tests should be attributed to the biological assumptions made in these tests.

## PART I – Identifying adaptive genes with high stringency

We first determined the distribution of the P values across genes. The MK test P values were obtained from the Fisher’s exact test on site count contingency tables. The likelihood ratio test was used to obtain PAML P values. The P value distributions are shown in the four panels of Fig. 2 for two taxa and two tests. The distribution is concentrated above P = 0.8 (the MK test for *Drosophila*) and P = 0.9 (the other panels). This concentration means that a very large percentage of genes show no detectable signal, partly because most genes experience too few changes to be statistically informative. Furthermore, the null model does not fully incorporate factors that can affect the test. For example, the polymorphism data may not reflect the complete removal of deleterious mutations and the strength of negative selection is often under-estimated^4,31,35^.

**Figure 2.**
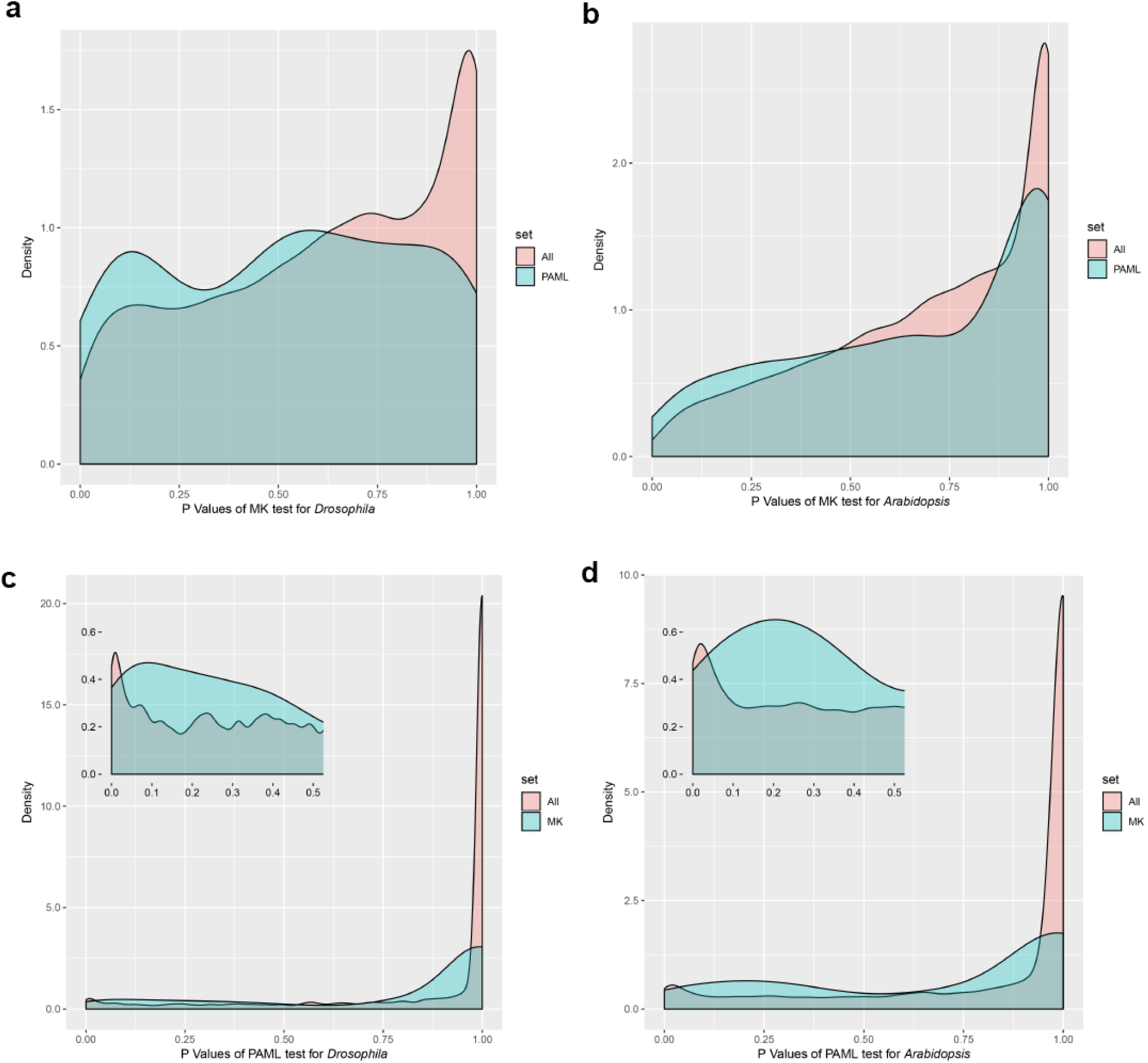
P value distributions of the MK and PAML test. **a** and **b**, P values of the MK test for *Drosophila* and *Arabidopsis*. The distribution for all genes is shown in red and the distribution for genes pre-filtered by the PAML test is shown in blue. **c** and **d**, P values of the PAML test. Results of genes pre-filtered by the MK test is shown in blue. These two panels are the mirror images of panels (a-b) with MK and PAML switched.

Fig. 2 suggests that, even if all genes evolve neutrally, far fewer than 5% of them would be detected as adaptive at the 5% cutoff. We therefore compare the observed P values from the MK and PAML tests against each other, rather than against the null model. In each panel of Fig. 2, one line represents the test results on all genes and the other is derived from loci that have been pre-filtered through the other test. In Fig. 2a-2b, genes pre-filtered through PAML have smaller P values in the MK test, reflected by the leftward shift in the P value distribution. The same is true in Fig. 2c-2d where pre-filtering by MK reduces the PAML test P values. The two tests are indeed correlated, but only weakly. This is also true in Fig. S1, where the branch-site model of PAML is used.

We now enumerate the overlap between the two tests by comparing the candidate adaptive genes with P < 0.05. Given the P value distributions shown in Fig. 2, these genes are merely the most likely candidates proposed by each test. Hence, significant overlaps would be mutual corroborations. For the “individual genes” analysis in *Drosophila*, we identified 186 from 5425 genes by the MK test and 145 genes by PAML, corresponding to 3.43% and 2.67% of the genome (see Table 1). The overlap between these two sets contains only nine genes. Although the observed overlap is higher than the expected 4.97 (P < 0.1, Fisher’s exact test), the overlap is too small to be biologically meaningful. The same pattern is true for *Arabidopsis*, in which 145 and 505 genes are called by these two tests but only 14 genes are called by both tests. Again, the observed overlap is significantly higher than the expected 5.55 (P < 0.01, Fisher’s exact test) but the actual overlap is minimal. A simple explanation for the non-overlap is a high false-negative rate. In other words, each test may have detected only a small fraction of the true adaptive genes.

**Table 1.** Proportion of adaptively evolving genes identified by two tests (P < 0.05)

| **Gene Category** | **MK** | **PAML** | **Expected overlap** | **Observed overlap** |
| --- | --- | --- | --- | --- |
| *Drosophila* |  |  |  |  |
| Individual genes | **3.43%**<br>(186/5425) | **2.67%**<br>(145/5425) | **0.09%**<br>(4.97/5425) | **0.17%**<br>(9/5425) |
| Supergenes<sup>a</sup> | **56.0%**<br>(112/200) | **18.0%**<br>(36/200) | **10.1%** | **10.0%**<br>(20/200) |
| Component genes<sup>b</sup> | **5.04%**<br>(158/3132) | **5.77%**<br>(60/1040) | **0.29%** | **0.48%**<br>(3/619) |
| *Arabidopsis* |  |  |  |  |
| Individual genes | **1.12%**<br>(145/12975) | **3.89%**<br>(505/12975) | **0.04%**<br>(5.55/12975) | **0.11%**<br>(14/12975) |
| Supergenes | **8.20%**<br>(41/500) | **25.6%**<br>(128/500) | **2.10%** | **2.00%**<br>(10/500) |
| Component genes | **3.63%**<br>(38/1048) | **7.44%**<br>(246/3306) | **0.27%** | **0.78%**<br>(2/258) |

<sup>a</sup> Supergenes are concatenations of 20-30 neighboring genes by physical location. See Table S1 for supergenes concatenated by gene function.
<sup>b</sup> Component genes are individual genes within supergenes that have passed the MK and/or PAML test.

### The analysis of supergenes and their component genes

False negatives should be common when analyzing individual genes. Since a gene on average harbors only a few substitutions, the power to reject the null model is often low. To augment the statistical power, we created artificial “supergenes” by merging 20 to 30 genes into a longer sequence. In the statistical sense, a supergene is like any standard gene that comprises a string of sites, each with a different adaptive value. Here, supergenes are either concatenations of neighboring genes (i.e., by physical location) or genes of the same ontology (by function). The merger would reduce false negatives due to low substitution numbers, but at the risk of diluting true adaptive signal. We present the results based on the concatenations of neighboring genes in Table 1. In *Drosophila* and *Arabidopsis*, 200 and 500 supergenes are created respectively. The results based on the merger by gene ontology are similar (See Table S1).

Our gene merger approach may create biases in the MK test, as pointed out before^4^. When the level of polymorphism is negatively correlated with the rate of nonsynonymous divergence across loci, false positives would be common in the merger. Hence, we used the modified MK test to infer positive selection in merged genes^4^. In *Drosophila*, 112 of the 200 supergenes reject the MK test null hypothesis at the 5% level, and 36 of the 200 significantly deviate from the PAML null (Table 1). The two tests detect far more adaptive supergenes than individual genes: 56% (MK) and 18% (PAML). What is perplexing being that the overlap between the two sets is random (10.0% observed vs. the expected 10.1%), as if the two tests are completely uncorrelated. In *Arabidopsis*, 8.2% of the 500 supergenes pass the MK test at the 5% level and 25.6% of supergenes reject the PAML null. The PAML test in *Arabidopsis* detects many more adaptive supergenes than the MK test, in the opposite direction of *Drosophila*. However, the overlap is also random with 2.0% observed vis-à-vis the expected 2.1%. In both taxa, the two tests appear uncorrelated at the level of supergenes.

Because gene merger might dilute the adaptive signal by mixing a few adaptively evolving genes with many other non-adaptive genes, we examined the component genes within each adaptive supergene. In *Drosophila*, the 112 supergenes passing the MK test contain 3132 component genes (Table 1), among which 158 genes are significant when tested individually. Likewise, 60 out of 1040 component genes are identified by PAML. Between the two subsets of genes (3132 and 1040), 619 genes are common and only three genes are significant by both tests. The 0.48% overlap of component genes is slightly higher than the expected 0.29%. The observations in *Arabidopsis* are given in the last row of Table 1. The overlap in component genes is also very low, at two of the 258 genes, or 0.78%. Clearly, the MK and PAML tests are uncorrelated by the standard statistical criteria, which are relaxed in the next section. Comparable analyses using the PAML branch-site model (Table S2) yield results similar to those in Table 1.

## PART II – Identifying weakly adaptive genes with low stringency

We note in Fig. 2 that genes yielding a P value of 0.25 by either test may be moderately informative about positive selection. Therefore, when carrying out the MK and PAML tests simultaneously, we set the cutoff in each test at P < 0.224. By doing so, the expected overlap would be 0.224^2^ = 5% if the two tests are completely uncorrelated. The results by this relaxed stringency are given in Table 2.

**Table 2.** Proportion of adaptively evolving genes identified by two tests (P^2^ < 0.05, i.e. P < 0.224)

|  | **MK** | **MK-PAML**<br>**overlap** | **PAML** | **Total** |
| --- | --- | --- | --- | --- |
| *Drosophila* |  |  |  |  |
| No. of genes | 824 | 91<sup>d</sup> | 353 | 5425 |
| Expected overlap | / | 53.6 | / | / |
| Proportion of adaptive changes by MK<sup>a</sup> | 0.69 | 0.67 | 0.32 | 0.26 |
| No. of adaptive sites per gene by MK (A1) | 14.98 | 19.94 | 5.71 | 2.84 |
| No. of adaptive sites per gene by PAML (A2)<sup>b</sup> | 10.93 | 14.65 | 9.27 | 5.71 |
| No. of adaptive sites per gene by PAML (A2')<sup>c</sup> | 3.19 | 8.62 | 6.24 | 1.79 |
| *Arabidopsis* |  |  |  |  |
| No. of genes | 1014 | 119<sup>e</sup> | 1172 | 12975 |
| Expected overlap | / | 91.6 | / | / |
| Proportion of adaptive changes by MK | 0.69 | 0.67 | 0.06 | 0.04 |
| No. of adaptive sites per gene by MK (A1) | 19.36 | 28.98 | 1.97 | 0.84 |
| No. of adaptive sites per gene by PAML (A2) | 14.24 | 20.36 | 12.74 | 10.15 |
| No. of adaptive sites per gene by PAML (A2') | 3.59 | 11.51 | 8.13 | 2.44 |

<sup>a</sup> Proportion of adaptive changes is done using Shapiro *et al.*'s method of correction<sup>4</sup>.
<sup>b</sup> A2 is based on PAML-M2a model.
<sup>c</sup> A2' is based on PAML-BEB model.
<sup>d</sup> $P < 10^{-7}$ by Fisher's exact test, given 53.6 as the expected value.
<sup>e</sup> $P < 0.002$ by Fisher's exact test, given 91.6 as the expected value.

The MK test identifies 824 and PAML 353 genes in *Drosophila*. These sets have 91 loci in common, whereas the expected overlap is 53.6 (P < 10^−7^, Fisher’s exact test). In *Arabidopsis*, the two tests yield 1014 and 1172 genes with an overlap of 119 genes, significantly higher than the expected number of 91.6 (P < 0.002, Fisher’s exact test). Hence, the joint call of adaptive genes accounts for 10.1% (119/1172) to 25.8% (91/353) of the loci identified by each single test. A gene identified by one test as adaptive has a 10% to 25% chance of being called adaptive by the other.

While the overlap between the two tests is at most modest, the performance of one test conditional on the pre-screen by the other indeed suggests some concordance. We first look at A1, the average number of adaptive sites per gene estimated using the MK test. A1 doubles from 2.84 to 5.71 when genes are pre-screened using PAML in *Drosophila* and increases from 14.98 to 19.94 in loci identified by both tests compared to just MK. The trend is even more pronounced in *Arabidopsis*: 0.84 to 1.97 and 19.36 to 28.98. Thus, the PAML screen can enhance the performance of the MK test.

The procedure is now applied in the reverse direction by pre-screening the genes with the MK test before subjecting them to the PAML test. The number of adaptive sites per gene can be calculated using two methods in PAML (A2 and A2’ in Table 2^37,39^; see Methods). Since the purpose is to compare PAML with MK, we use the A2 numbers, which are closer to A1 from the MK test. The qualitative conclusion, nevertheless, is not affected much by the choice of model. The number of A2 sites increases from 5.71 to 10.93 after MK pre-screening in *Drosophila* (Table 2) and from 9.27 to 14.65 when focusing on the loci identified by both PAML and the MK test, compared to PAML alone. The same trend is observed in *Arabidopsis* (Table 2): an increase from 10.15 to 14.24 after MK test pre-screening and 12.74 for PAML only vs 20.36 for genes identified by both tests. Again, a pre-screen by MK helps PAML performance.

The results are similar when we use the PAML branch-site model rather than the site model (Table S3). It is clear that the MK and PAML tests are correlated but the correlation is weak. In other words, when one test detects a strong adaptive signal in a gene, the other test would often find a signal in the same gene, albeit a much weaker one.

## Discussion

This may be the first study that compares the MK and PAML tests for the inferences of adaptive evolution on the same set of genes along the same phylogenetic branch. Many previous studies have also employed the two tests, albeit for different purposes^25,40–44^ (see Supplementary Information). It is surprising that the two widely used tests are so poorly concordant in detecting adaptively evolving genes. We examine possible explanations below.

### i) One of the two tests has high false-positive and false-negative rates

By this explanation, one of the two tests is entirely unreliable. However, the two tests are comparable in performance. When PAML is done on genes selected by the MK test, the subset of genes yields much stronger signal than the full set. This is also true when the MK test is done on PAML-selected genes. Apparently, these two tests yield comparable quantitative results.

### ii) Both tests have high false negative rates due to low statistical powers

This may be the most obvious explanation. If the fraction of genes driven by positive selection is high and the power of detection is low by both tests, the overlap in the gene sets identified may indeed be low. False negatives of this kind could be a consequence of the “nearly neutral” evolution proposed by Ohta (1992)^45^ and Ohta and Gillespie (1996)^29^. However, false negatives are unlikely to be correct. For example, the fraction of *Drosophila* supergenes yielding adaptive signals is 0.560 and 0.180, respectively, for MK and PAML (Table 1). Since the observed overlap at 0.100 is exactly the same as the overlap between random picks (0.101), true adaptive genes have to account for close to 100% of the total, if the explanation is correct.

### iii) Both tests have some false negatives due to different biases: both are, at least partially, correct

The two tests may be complementary. In other words, their results would only partially overlap even when all are correct with no false positives. A main reason is that the detection of positive selection is influenced by the strength of negative selection operative on the same genes. This explanation may diverge from some of the tenets of molecular evolution and is presented in the Supplement (see also Chen *et al.*^46^).

### iv) False positives caused by fluctuating negative selection: both tests will require modifications

In order to detect positive selection, the intensity of negative selection is assumed to be constant in both MK and PAML. Otherwise, it would be impossible to disentangle positive and negative selection from Eq. (1). However, it has been shown recently that this constancy assumption is violated in all taxa analyzed^1^. Since false positives in the two tests are influenced in different ways by changing negative selection, genes falsely identified as adaptively evolving would be correspondingly different.

The MK test assumes that the negative selective pressure in the extant polymorphism accurately reflects the average of the earlier periods. Fig. 1 of Chen *et al.*^1^ clearly invalidates this assumption. The same figure also invalidates the assumption of PAML whereby negative selection is assumed constant in the phylogenetic branches of interest. In fact, even a casual glance of the Arabidopsis data would reveal the flaw of the constancy assumption. Between *A. thaliana* and *A. lyrata*, the Pa/Ps ratio is 0.167 and 0.271 but the Ka/Ks ratio is 0.215 (Table S2 of Chen *et al.*^1^). Clearly, the strength of negative selection has changed in this short time span. Furthermore, by the MK test, one would reach opposite conclusions depending on the polymorphism data chosen.

## Conclusion

In the search for the signals of positive selection, the extensive literature of the last two decades has neglected the continual changes in negative selection, which often overwhelm the adaptive signals. To gauge the strength of negative selection will entail the use of polymorphism data from multiple species^1^. Finally, an expanded framework that permits the simultaneous analyses of positive and negative selection will be necessary^46–48^.

## Methods

### DNA sequence data

Pre-aligned unique *Drosophila* transcript sequences were downloaded from Flybase^24^ (http://flybase.org). We collected 8560 FASTA alignments of five species (*D. melanogaster, D. simulans, D. sechellia, D. yakuba*, *and D. erecta*, Fig. 1a). Genome-wide *D. melanogaster* polymorphism data were obtained from the *Drosophila* Population Genomics Project Phase 2^49^. Genes with high divergence rates, apparently caused by misalignment, were discarded. Only genes with more than 40 codons and 10 samples of polymorphism data were used. The final dataset contains 5245 genes with an average of 50 samples of polymorphism data.

The DNA sequences of *A. thaliana* and *A. lyrata* were obtained from the Phytozome database^50^. Genome-wide *A. thaliana* polymorphism data were obtained from the 1001 Genomes Project^51^. We also obtained DNA sequences from *Capsella grandiflora*, *C. rubella*, and *Boechera stricta* (Fig. 1b) for analyses using the PAML program^27^. We began with 14953 alignments of the five species and then filtered the data as we did for *Drosophila*. Only genes with more than 300 samples of polymorphism data were collected. The final dataset consists of 12975 genes.

### Supergene construction

To overcome statistical limitations, we created artificial supergenes by merging genes into longer sequences. We used two concatenation approaches: by physical location and by ontology. The first method involved merging 20 to 30 nearby genes residing on the same chromosome. This resulted in 200 *Drosophila* and 500 *Arabidopsis* supergenes. To apply the ontology approach, we first identified GO (gene ontology) term(s) for each gene. To ensure that every gene was present in only one supergene, we sorted GO terms by the number of genes they comprised and checked the component genes in each supergene. If a gene was previously included in a set, it was not merged again. GO terms with fewer than eight genes in *Drosophila* and 10 in *Arabidopsis* were discarded. The final set comprised 184 *Drosophila* and 454 *Arabidopsis* supergenes.

### The McDonald-Kreitman (MK) test

Let A and S be the number of nonsynonymous and synonymous changes per gene (or per genome). In Fig. 1c-1d, A/S ratio is given for the number of polymorphic changes at a defined frequency range. (The frequencies were inferred by the free-ratio model of the PAML site module)^26,27^. The A/S ratio becomes lower when the mutant frequency becomes higher. Apparently, the A/S ratio at the low frequency range is boosted by deleterious mutations that have not been removed by negative selection. To avoid the confounding effect of negative selection on the MK test, we only used common mutations with derived allele frequencies larger than 0.2, as is done previously^4,52^. Note that the A/S ratio reaches a steady level at around 0.2 in Fig. 1c-1d.

Now, we let A and S designate the total number of *common* polymorphic mutations (frequency > 0.2) in the MK test. The corresponding numbers of changes between species are designated A’ and S’. These four numbers are gathered in a 2×2 contingency table. Fractions of amino acid substitutions which are adaptive (α) can be estimated as α = 1 − (S’ A) / (A’ S). We used Fisher’s exact test on 2×2 contingency tables to estimate statistical significance. Shapiro *et al.* pointed out the possibility of false positives when the MK test is applied across genes and proposed a procedure to correct the bias^4^. Hence, we used it in the calculations.

Ideally, the number of A and S polymorphic sites should reflect only neutral variation. However, as can be seen in Fig. 1c-1d, A often includes low-frequency deleterious mutations and, perhaps, high-frequency advantageous mutations^2,4,5,20^. The inclusion of both kinds of mutations would bias the polymorphic A/S ratio upward, hence reducing the excess of A’/S’ over A/S and compromising the power of the MK test. Various solutions have been proposed^3,4,31,35^ to more accurately measure the polymorphic A’s. These methods are mostly ad hoc in nature. Sawyer and Hartl^17^ propose a more robust approach to this problem by directly estimating the intensity of negative selection. While the theory outpaced the data at that time, the approach is feasible now given the large amount of polymorphism data.

If the distribution of the strength of negative selection is known^17^, the neutral A/S ratio as reflected in the polymorphism can be accurately estimated. While the estimation of positive selection is indeed different from the conventional numbers, the MK results obtained by various procedures do show the same qualitative pattern of limited overlap with the PAML test. The overall patterns suggest that the discordance between the MK and PAML tests is biological, rather than technical, as presented in Discussion.

### The PAML test

We used both the site model and the branch-site model in PAML. The site model, allowing the ω ratio (dN/dS) to vary among sites (codons or amino acids in protein), detected positive selection across the five chosen species. A likelihood ratio test (LRT) was used to compare the alternative model M2a (selection model allowing an additional category of positively selected sites with ω > 1 by setting: model = 0, NSsites = 2, fix_omega = 0, omega = 2) with the null model M1a (neutral model allowing only two categories of sites with ω < 1 and ω = 1 by setting: model = 0, NSsites = 1, fix_omega = 0, omega = 2). Significance was determined using the chi-squared test (df = 2).

The branch-site model, allowing dN/dS to vary both among sites and across lineages, was used to detect positive selection along specified branches. We compared the likelihood of the alternative model A (positive selection, model = 2, NSsites = 2, fix_omega = 0), to the null model A1 (model = 2, NSsites = 2, fix_omega = 1, omega = 1). *D. melanogaster* and *A. thaliana* were designated as the foreground branches for the test. Significance was calculated using LRT as above. The site model results are presented in the text and the branch-site results are given in Supplementary Information.

In the analysis of both models, we also employed Bayes empirical Bayes (BEB)^9^ estimates, which are available for calculating the posterior probabilities for site classes and can be used to identify sites under positive selection if the likelihood ratio test is significant.

## Acknowledgements

We thank Daniel Hartl, Ziheng Yang, Adam Eyre-Walker, Dan Graur, Justin Fay, Bruce Rannala and Matthew Hahn for their helpful discussions. This study was supported by the National Natural Science Foundation of China (31830005 and 31971540); the National Key Research and Development Plan (2017FY100705); the 985 Project (33000-18831107); and Guangdong Basic and Applied Basic Research Foundation (2019A1515010752).

