## Supplementary Information for "Two decades of suspect evidence for adaptive DNA-sequence evolution - Failure in consistent detection of positive selection"

**This file includes:**

Supplementary Note

Supplementary references 53-59

Tables S1-S3

Figures. S1-S3

### Supplementary Note

#### Previous studies that employed MK and PAML tests

Some previous studies did use both MK test and PAML, but their purposes were varied. They can be roughly grouped into several categories. The first category used both of their results to make a conclusion but did not discuss too much about their inconsistencies. For instance, Luisi *et al.* studied how positive selection affects across parts of molecular networks using comparative genomics and population genetics approaches on human and related species' genomes[43]. This article concluded that relationship between centrality and the impact of adaptive evolution highly depends on the mode of positive selection and/or the evolutionary time-scale. Feldmeyer *et al.* used three methods (MK test, PAML branch model and TreeSAAP) to identify positively selected genes involved in species divergence on orthologous mt-genome transcripts of four closely related European Radix species[44]. In total 134 genes were identified as positively selected by at least one of the three methods. Of these, 116 were identified by a single method only, and 5 genes by all three methods. Wong *et al*. proposed that seminal fluid proteases are likely targets of selection due to their demonstrated or potential roles in between-sex interactions and immune processes. The gene pool is limited to five predicted protease-encoding *Acp* loci to test this hypothesis, and several within- and between-species sequence methods were used, including the MK test and PAML[53]. One gene was identified by the MK test whereas PAML found another two. Biswas *et al*. describe the identification of targets of adaptive evolution[40]. They reviewed previous studies using different methods to detect positive selection based on polymorphisms within species, as well as polymorphism within and divergence between species. Some inconsistencies in these results have been observed but this paper offers three simple explanations: 1) different studies are probably detecting different selective events; 2) even for tests that should detect similar types of selective events, low statistical power further decreases the probability of overlap; 3) most studies report only the most significant results.

The second category use one method to detect adaptive evolution and adopt the idea from the other as a supplement. For example, Nielsen, R. *et al.* was comparing 13,731 annotated genes from humans to their chimpanzee orthologs to identify genes that show evidence of positive selection[25]. They identified 50 genes with the highest likelihood ratio. PAML was used to validate those genes if the elevated dN/dS was in the human lineage, chimpanzee lineage or both lineages. Population genetic analysis of nonsynonymous polymorphisms of those 50 genes showed evidence for an excess of high frequency alleles, providing additional support for positive selection. Gayà-Vidal *et al*. used the MK test $\alpha$ for estimating the amount of positive selection in the human lineage[42]. It considered several factors that could have an impact on $\alpha$: 1) high derived allele frequency (up to 60%) will avoid the impact of slightly deleterious mutations and thus elevate $\alpha$; 2) Human accelerated genes (dN/dS accelerate in human branch) and human BS test genes (genes detected by PAML branch-site model) would elevate the $\alpha$ but the effect of the former one was better; 3) Pre-screening for rapidly evolving genes performed better than using mammalian-specific genes.

The third category applied the ideas from either (or both) of these two methods to generate a new tool and make new discoveries. Welch *et al*. introduced an improvement for the MK estimate $\alpha$ because the conventional way of doing so might be subjected to biases such as reflecting demographic changes rather than adaptive substitution; Using between-locus variation yielded contradictory results; Estimation from the same model organisms have also varied widely[41]. Welch *et al.* used three sets of dS and dN measurements: the total divergence between *D. yakuba* and *D. simulans*, the total divergence between *D. melanogaster* and *D. simulans,* and the divergence along the *D. simulans* lineage alone, estimated using PAML.

The last category is pure theoretical comparison based on simulations. Zhai *et al*. investigated the statistical power of three neutrality tests for comparative data[54]: the HKA test[55], the MK test[13], and the dN/dS likelihood ratio test[15,39], along with two tests based on population genetic data. Using forward simulations, this paper shows that: 1) the most important role of the MK test in population genetics might perhaps be to test for negative selection, whereas other tests should be used to detect positive selection. 2) Under the assumption of a fixed-position model, the dN/dS ratio has more power to detect recurrent positive selection than any of the tests which use population genetic data. 3) If more species are included and/or if the divergence time is longer, the power of dN/dS ratio tests increases.

**Supplementary note of theoretical background**

Kimura[30] has showed the fixation probability $f_1$ of an advantage mutant is

$$f_1 = \frac{1-e^{-S_1}}{1-e^{-2NS_1}}$$

where N is population size of a diploid population. The relative fitness of three alleles are $1+S_1$, $1+S_1/2$ and 1 ($S_1>0$).

Hence, the fixation probability $f_2$ of a deleterious mutant could be derived as

$$f_2 = \frac{1-e^{\frac{-S_2}{1+S_2}}}{1-e^{\frac{-2NS_2}{1+S_2}}}$$

The relative fitness of three alleles are $1+S_2$, $1+S_2/2$ and 1 ($S_2<0$). Given $|S_2| << 1$, the above formula may be simplified as

$$f_2 \approx \frac{1-e^{-S_2}}{1-e^{-2NS_2}}$$

Hence, $f \approx (1 - e^{-s}) / (1 - e^{-2Ns})$ is the fixation probability of a mutation with a selective coefficient s that can be > 0 (denoted by $s_1$) or < 0 ($s_2$) and half the value in heterozygotes.

On the other hand, the fixation probability f of a neutral mutant is 1/2N[28,56]. Then, Ka/Ks = f / (1/2N) = 2Nf. The Ka/Ks of total sequences could be the weighted sum of the three parts (neutral, advantage and deleterious) as shown in Eq. (1) in the end.

**The results of branch-site model**

The results of using branch-site model of PAML are displayed in Fig. S1 and Table S2-S3. The result in Fig. S1 is similar to Fig. 2 that used site model of PAML. Likewise, similar to Table 1, Table S2 displayed the overlaps between PAML and MK tests with the threshold $P < 0.05$. In *Drosophila* species, 293 and 186 genes (5.40% and 3.43%) were called by PAML (branch-site model) and MK test, respectively. Only 19 genes are called by both tests, though it is significantly higher than the expected 10.05. Among the 12975 genes in *Arabidopsis*, only a few genes are overlap between two tests (21 genes, 0.16% of the genome). Results of the lower stringency ($P < 0.224$) similar to Table 2 are given in Table S3.

To reduce false negatives, 20-30 neighboring genes were merged into "supergene". In *Drosophila*, 16 out of the 200 supergenes are detected by both tests (Table S2), but their overlap is still almost at random (8.00%, slightly less than the expected 10.08%). In *Arabidopsis,* the two tests also appear to be weakly correlated (observed 2.20% vs. the expected 3.87%). In addition, similar to the result of site model (see Table 1), the overlap analysis of component genes did not give a satisfactory result.

**Discussion iii) Both tests have some false negatives due to different biases: both are, at least partially, correct**

Under negative selection, the fixation probability decreases rapidly when the strength ($Ns_2$) increases (Fig. S2a). This strength reflects the functional importance (x) of the gene, or the part of the gene hit by mutations. For example, mutations in the heme-pocket of hemoglobin would have a larger x whereas those in the backbone of the same protein would have a smaller x. This functional attribute may influence the performance of both MK and PAML.

In particular, PAML does not filter out signals of negative selection; hence, mutations associated with a smaller x could be more readily pushed above Ka/Ks > 1 by positive selection (see Fig. S2b) whereas genes under stringent constraints may require very strong positive selection to appear adaptive. Fig. S3 indeed shows that genes with a higher polymorphism A/S (non-synonymous / non-synonymous changes) ratio (i.e., weaker negative selection) tend to show a stronger signal of positive selection. In contrast, the MK test detects positive selection by comparing the divergence A/S ratio with the polymorphism, Pa/Ps. Hence, a smaller Pa/Ps that reflects stronger negative selection would permit better detection of positive selection by the MK test (see Fig. S2b).

The opposite patterns of Fig. S2b provide a rationale for the "both are right" hypothesis, but the low overlap between MK and PAML may still depend on the abundance of adaptive mutations as a function of x (Fig. S2c vs S2d). Fig. S2c illustrates the conventional view whereby positive selection tends to act on genes where negative selection is weaker (i.e., smaller x). This convention is accepted by both the selectionism and neutralism schools. According to the neutral theory, negative selection is weaker when a gene, or the mutation, is functionally less important (Rules 2 and 3 of neutrality; see p. 103 of Kimura, 1983)[57]. In parallel, in Fisher's geometric model of adaptive evolution[58,59], positive selection favors mutations of smaller functional changes. In the scheme of Fig. S2c, the MK and PAML results overlap substantially, thus failing to support the “both are right” explanation for the reported pattern in this study.

In contrast to Fig. S2c, Fig. S2d presents an opposite scenario by postulating that both positive and negative selection should become stronger as x increases. After all, larger functional differences may be more “discernible” to selection, regardless of the direction of selection. The overlaps between MK and PAML would indeed be much smaller in Fig. S2d (47% of the detected genes overlap) than in Fig. S2c (87% overlap). Although the theoretical overlap shown in Fig. S2d is still far larger than the observed value, the trend is instructive. The key message is that the distribution in Fig. S2d, which is far more likely to support the complementarity hypothesis, is not compatible with the conventional view (Fig. S2c). While some recent efforts have challenged this conventional view[46], it should be noted that the overlaps reported in Tables 1 and 2 are too small to dispel the doubt of high false positives.

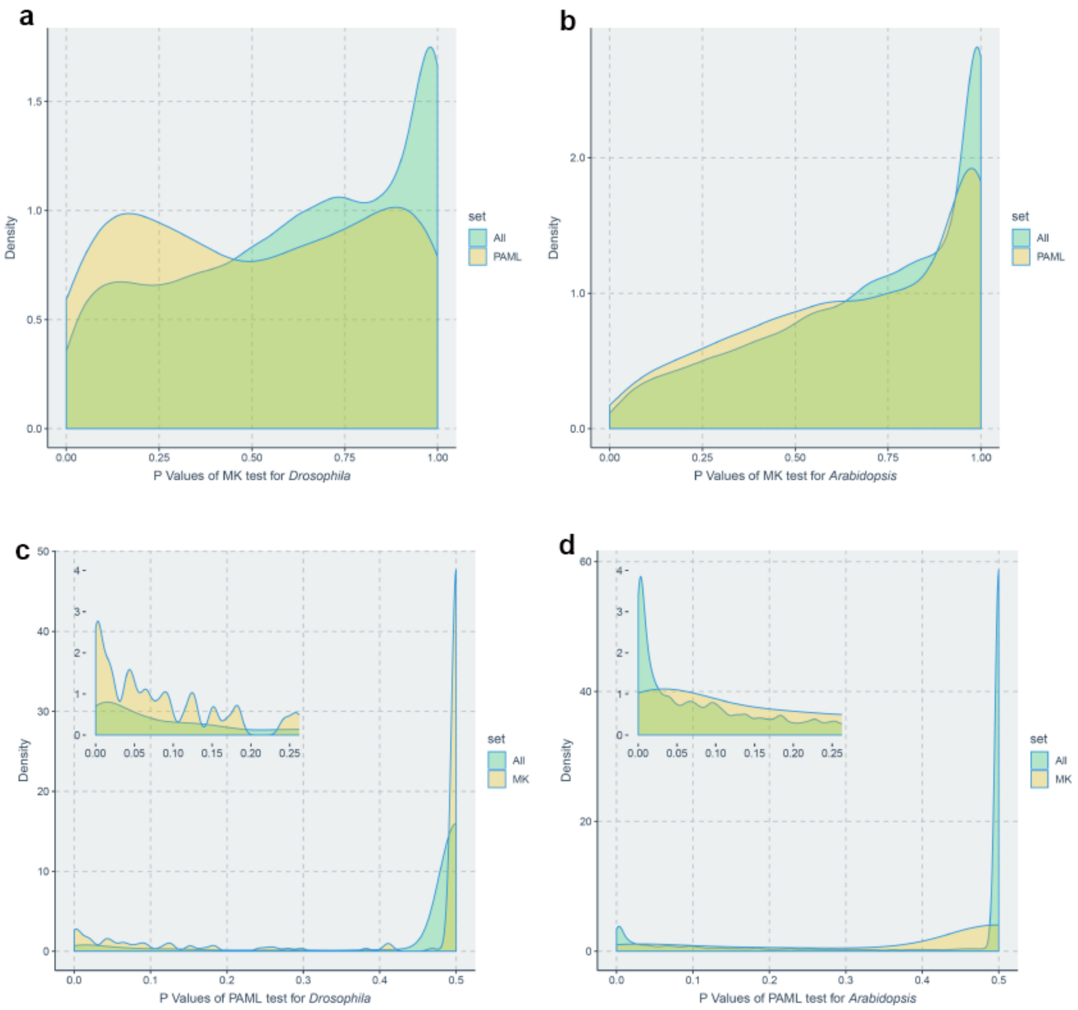


**Figure S1. P value distributions of MK and PAML tests (the branch-site model).** In panels (a) and (b), the P values of the MK test for all genes are shown in green, and that for genes selected by PAML test are shown in yellow. In panels (c) and (d), the green distributions represent the P values of the PAML test for all genes, and the yellow distributions the genes selected by MK test.

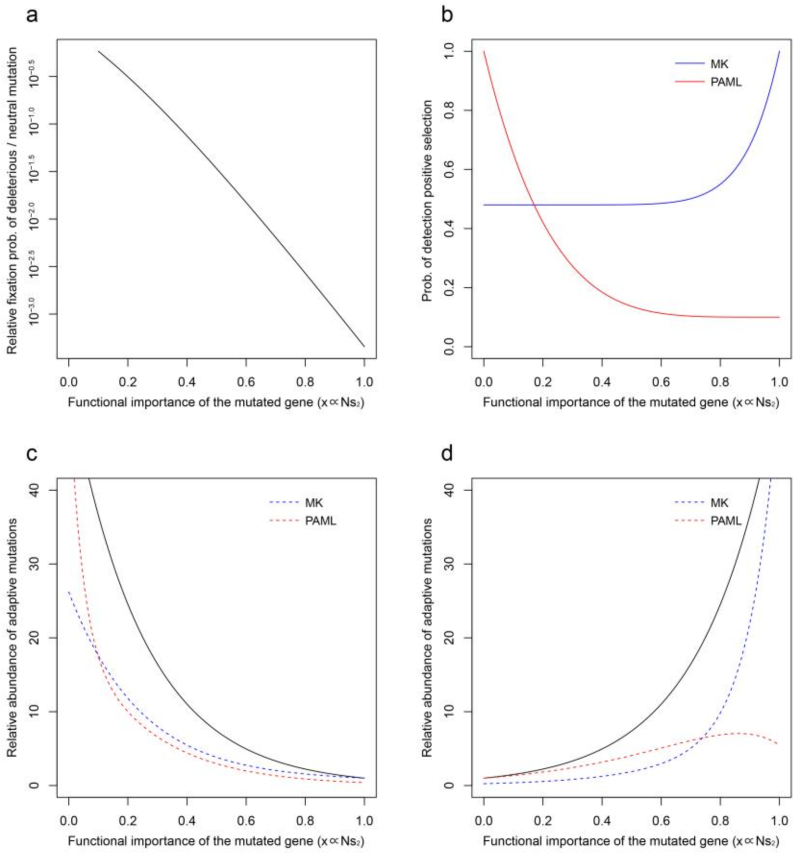


.

**Figure S2. Models for detecting positive selection under different strength of negative selection.** x, an arbitrary scale for the functional importance of the mutated gene, determines the strength of negative selection. **a**, The fixation probability of deleterious mutations based on Eq. (2). **b**, The probability of detecting adaptive mutations ($P^+$) by MK or PAML. For MK, the blue line follows the equation $P_{mk}(x) = (1\text{-}c) + cx^i$ and, for PAML, the red line follows the equation $P_{paml}(x) = (1\text{-}c) + c(1\text{-}x)^i$. The assumption is that MK and PAML has the maximal efficiency at x = 1 and x = 0, respectively (see text). **c**, The relative number of adaptive mutations, $M = e^{4(1-x)}$, as a function of x (black line). The blue line, the product of $M\ P^+ = M\ [(1\text{-}c) + cx^i]$, denotes the relative number of genes detected by MK. The red line, $M\ [(1\text{-}c) + c(1\text{-}x)^i]$, denotes the corresponding number detected by PAML. It is assumed that both tests detect 50% of the adaptive genes. Note that the high detection rate is based on the number of adaptively-evolving genes whereas the empirical observations in Table 1 are based on the number of all genes. The overlap in detection [ = Min ($P_{mk}(x)$, $P_{paml}(x)$)] between the two tests is 87% of the detected genes. **d**, same as c but $M = e^{4x}$. The MK test still detects 50% but the PAML test can only detect 30% due to the higher density of genes with a larger x. The overlap accounts for 44% of the MK detection. In short, the overlap between the two tests depends on the distribution of adaptive mutations as a function of x; hence, the overlap in Fig. S2d can be much smaller than that in Fig. S2c.

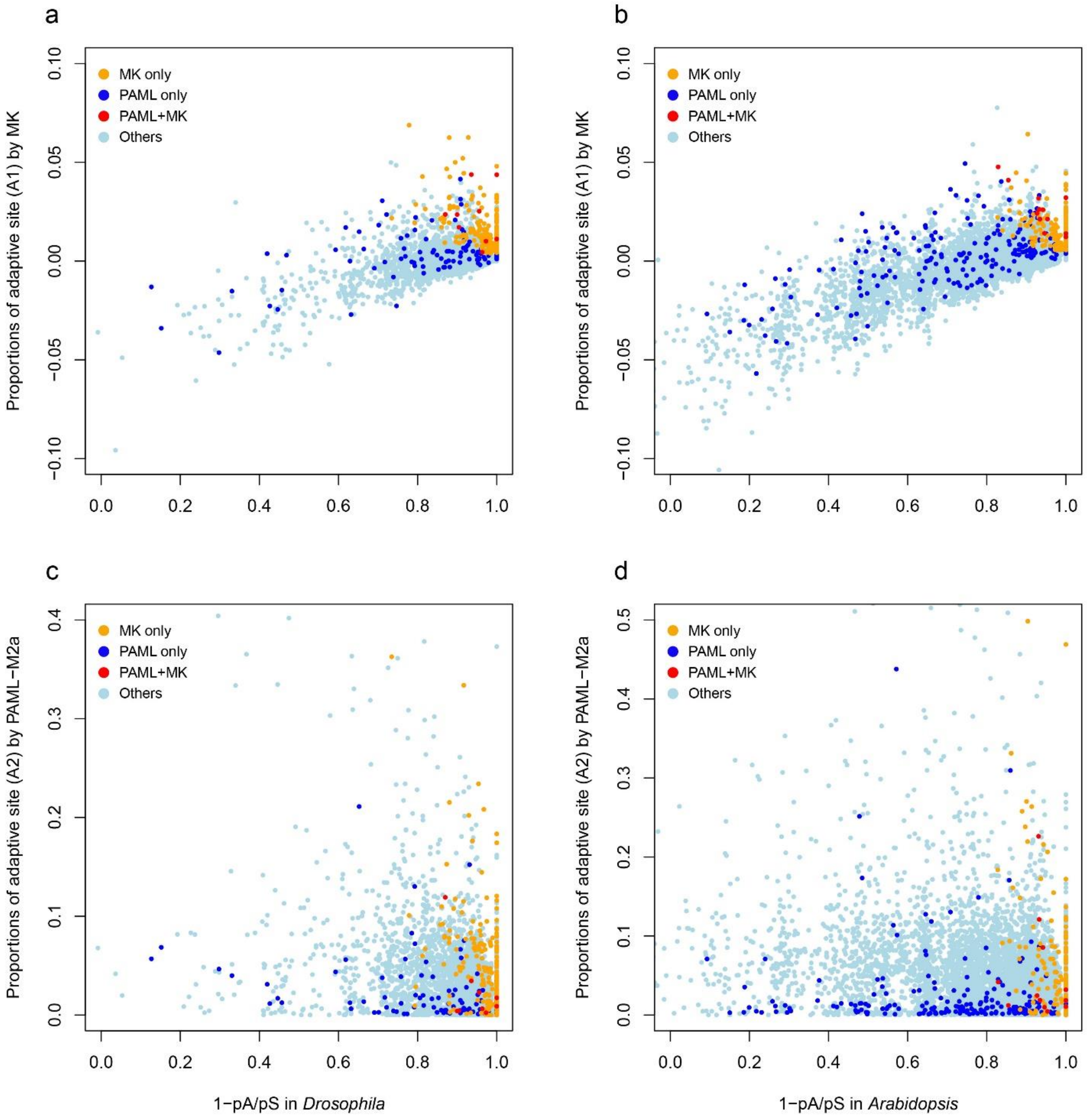


**Figure S3. Scatter plots of the proportion of average adaptive sites identified by MK and PAML tests (the site model).** The orange dots represent genes chosen by the MK test as candidate genes under positive selection, but not chosen by the PAML test. Genes with more than eight polymorphisms are shown. Genes identified by the PAML but not the MK test, are depicted by dark blue dots. Red dots are genes called by both tests. The remaining genes are represented by light blue dots.

**Table S1. Proportion of adaptively evolving genes identified by two tests (P < 0.05)**
**(Same as Table 1 but genes were merged into supergenes by ontology)**

| Gene Category | MK | PAML (site model) | Expected overlap | Observed overlap |
|---|---|---|---|---|
| ***Drosophila*** | | | | |
| Supergenes[a] | **31.52%** (58/184) | **14.67%** (27/184) | **4.62%** | **8.70%** (16/184) |
| Component genes[b] | **6.01%** (51/849) | **7.54%** (36/477) | **0.45%** | **0.00%** (0/306) |
| ***Arabidopsis*** | | | | |
| Supergenes[a] | **10.57%** (48/454) | **19.38%** (88/454) | **2.05%** | **2.42%** (11/454) |
| Component genes[b] | **4.46%** (45/1008) | **7.19%** (184/2556) | **0.32%** | **0.29%** (1/341) |

[a] Supergenes are the concatenations of genes of the same ontology.

[b] Component genes are individual genes within supergenes that have passed the MK and/or PAML tests.

**Table S2. Proportion of adaptively evolving genes identified by two tests (P < 0.05)**

**(Same as Table 1 but using the PAML branch-site model)**

| **Gene Category** | **MK** | **PAML (branch -site)** | **Expected overlap** | **Observed overlap** |
|---|---|---|---|---|
| ***Drosophila*** | | | | |
| Individual genes | **3.43%** (186/5425) | **5.40%** (293/5425) | **0.19%** (10.05/5425) | **0.35%** (19/5425) |
| Supergenes[a] | **56.00%** (112/200) | **18.00%** (36/200) | **10.08%** | **8.00%** (16/200) |
| Component genes[b] | **5.04%** (158/3132) | **9.41%** (92/978) | **0.47%** | **1.76%** (8/455) |
| ***Arabidopsis*** | | | | |
| Individual genes | **1.12%** (145/12975) | **10.02%** (1300/12975) | **0.12%** (14.53/12975) | **0.16%** (21/12975) |
| Supergenes | **8.20%** (41/500) | **47.20%** (236/500) | **3.87%** | **2.20%** (11/500) |
| Component genes | **3.62%** (38/1048) | **12.24%** (750/6129) | **0.44%** | **1.36%** (4/295) |

[a] Supergenes are concatenations of 20-30 neighboring genes by physical location.

[b] Component genes are individual genes within supergenes that have passed the MK and/or PAML tests.

**Table S3. Proportion of adaptively evolving sites identified by two tests ($P^2 < 0.05$, i.e. $P < 0.224$)**

**(Same as Table 2 but using the PAML branch-site model)**

| | **MK** | **MK-PAML overlap** | **PAML (branch-site)** | **Total** |
|---|---|---|---|---|
| ***Drosophila*** | | | | |
| No. of genes | 824 | 127[d] | 530 | 5425 |
| Expected overlap | / | 80.50 | / | / |
| Proportion of adaptive changes by MK[a] | 0.69 | 0.65 | 0.31 | 0.26 |
| No. of adaptive sites per gene by MK (A1) | 14.98 | 20.22 | 6.24 | 2.84 |
| No. of adaptive sites per gene by PAML (A2)[b] | 8.33 | 23.96 | 22.40 | 5.02 |
| No. of adaptive sites per gene by PAML (A2')[c] | 1.95 | 9.69 | 9.53 | 1.23 |
| ***Arabidopsis*** | | | | |
| No. of genes | 1014 | 233[e] | 1172 | 12975 |
| Expected overlap | / | 193.89 | / | / |
| Proportion of adaptive changes by MK | 0.69 | 0.69 | 0.06 | 0.04 |
| No. of adaptive sites per gene by MK (A1) | 19.36 | 24.07 | 1.50 | 0.84 |
| No. of adaptive sites per gene by PAML (A2) | 4.48 | 9.94 | 7.78 | 3.33 |
| No. of adaptive sites per gene by PAML (A2') | 2.72 | 9.00 | 7.69 | 2.06 |

[a] Proportion of adaptive changes is done using Shapiro *et al.*'s method of correction[4].

[b] A2 is based on PAML-M2a model.

[c] A2' is based on PAML-BEB model.

[d] $P < 10^{-7}$ by Fisher's exact test, given 80.5 as the expected value.

[e] $P < 10^{-10}$ by Fisher's exact test, given 193.9 as the expected value.
